# Computational Drug Recommendation Approaches toward Safe Polypharmacy

**DOI:** 10.1101/518415

**Authors:** Wen-Hao Chiang, Xia Ning

**Affiliations:** Department of Computer & Information Science, Indiana University - Purdue University Indianapolis, Indianapolis, IN, USA; Department of Biomedical Informatics, The Ohio State University, Columbus, OH, USA

## Abstract

Adverse drug reactions (ADRs) induced from high-order drug-drug interactions (DDIs) due to polypharmacy - simultaneous use of multiple drugs - represent a significant public health problem. Unfortunately, computational efforts to facilitate decision making for safe polypharmacy, particularly to assist safe multi-drug prescribing, are lacking. We formally formulate the to-avoid and safe drug recommendation problems for multi-drug prescriptions. We investigate preliminary computational approaches to tackling these problems, utilizing a minimum set of available prescription data from a large population, and demonstrate their potentials in assisting safe-polypharmacy decision making once richer data (e.g., electronic medical records, omics data and pathology data) are available. We develop a joint model with a recommendation component and an ADR label prediction component to conduct to-avoid and safe drug recommendation. We also develop real drug-drug interaction datasets and corresponding evaluation protocols to facilitate future computational research on safe polypharmacy.

## Introduction

Adverse drug reactions (ADRs) induced from high-order drug-drug interactions (DDIs) due to polypharmacy - simultaneous use of multiple drugs - represent a significant public health problem. ADRs refer to undesired or harmful reactions due to drug administration. One of the major causes for ADRs is DDIs that happen when the pharmacological effects of a drug are altered by the actions of other drugs, leading to unpredictable clinical consequences. The increasing popularity of polypharmacy continues to expose a significant and growing portion of the population to unknown or poorly understood DDIs and associated ADRs. The National Health and Nutrition Examination Survey^1^ reports that more than 76% of the elderly Americans take two or more drugs every day. Another study^2^ estimates that about 29.4% of elderly American patients take six or more drugs every day.

Current research on DDIs and their associated ADRs is primarily focused on mining and detecting DDIs for knowledge discovery^2, 3^. Applying the knowledge in practice for preventive, proactive and person-centered healthcare also requires predictive power to deal with unknown DDIs, and to provide evidence-based suggestions to facilitate future drug prescribing. As wet-lab based experimental validation scheme for DDI study still falls behind due to its low throughput and lack of scalability, but meanwhile huge amounts of electronic medical record data become increasingly available, data-driven computational methodologies appear very appealing to tackle DDI and ADR problems. Unfortunately, computational efforts to facilitate future polypharmacy, particularly to assist safe multi-drug prescribing, are still in their infancy. In this manuscript, we present a computational approach toward the goal of facilitating future safe multi-drug prescribing. Here we use the term “prescription” to represent a set of drugs that have been taken together, even though there could be non-prescription drugs. Thus, we tackle the following *to-avoid drug recommendation* problem and *safe drug recommendation* problem.

### Definition 1. To-Avoid Drug Recommendation

*given the multiple drugs in a prescription, recommend a short, ranked list of drugs that should be avoided taking together with the prescription in order to avoid a particular ADR.*

### Definition 2. Safe Drug Recommendation

*given the multiple drugs in a prescription, recommend a short, ranked list of safe drugs that, if taken together with the prescription, are not likely to induce a particular ADR.*

We need to clarify and emphasize the following three aspects related to the recommendation problems and our approaches. First, in this pilot study, we only consider one ADR and thus drug safety is only considered with respect to that particular ADR. The one-ADR condition is over-simplified compared to the real scenarios, in which very often multiple ADRs will occur simultaneously. However, this simplification can enable well-calibrated comparison and evaluation of the prospective approaches that will be developed, and also can serve as a baseline upon which multi-ADR cases can be more easily tackled (e.g., as combinations of multiple, single ADRs, with correlation well addressed). Second, in this pilot study, we only use drug prescription data (i.e., which drugs are prescribed together) and their associated, single ADR, which is calculated based on a large population, in computational models. To truly enable precision medicine and safe polypharmacy, more and comprehensive data, such as electronic medical records, omics data, pathology data, etc., should be all integrated and utilized. However, our methods do not aim to fully solve the safe-polypharmacy problem, which is highly non-trivial and also requires close collaborations from clinicians, pharmacists, biologists, pathologists, etc. Instead, we aim to pioneer the development and test of computing power that can be leveraged to better solve the problem. Particularly, our methods use prescription data from a large population, which could be the most common, easy-to-access data type, and thus do not necessarily limit their potentials in real applications. In the end, our methods recommend individual drugs, although sorted, that should be avoided or safe with respect to an ADR. This is also likely that multiple drugs should be recommended as a composite, and/or the recommendations should take into considerations other factors such as efficacies and costs of the new drugs, etc., and thus multi-criteria recommendation. We cannot claim that our methods can satisfy all requirements in real practice. However, our methods can serve as a baseline upon which other comprehensive models can be developed (e.g., multi-criteria recommendation can be decomposed into multiple, single-criterion recommendations). To the best of our knowledge, this is still the first work to formally formulate the above problems and to provide a computational solution framework.

The two recommendation problems are significant particularly for healthcare practice. They can enable evidence-based suggestions to help polypharmacy decision making, and induce novel hypotheses on new high-order DDIs and associated ADRs. Our contributions to solving the two new drug recommendation problems are summarized below.

- We formally define the drug recommendation problems.
- We developed a joint sparse linear recommendation and logistic regression model (`SLR`), with a drug recommendation component and an ADR label prediction component (Section Joint `SLIM` and `LogR` Model: `SLR`) to solve the problems. The recommendation component captures drug co-prescription patterns among ADR and non-ADR inducing prescriptions, respectively, and uses such patterns to recommend drugs. The ADR label prediction component learns and predicts the ADR probabilities from drugs in prescriptions. These two components are learned concurrently in `SLR` so that the recommended drugs are more likely to introduce expected ADR labels.
- We developed a protocol to mine high-order DDIs and their associated ADRs, and provided a DDI dataset to the public (will be publicly available upon the acceptance of this manuscript) (Section Materials).
- We developed new evaluation protocols and evaluation metrics to evaluate the performance of drug recommendation (will be publicly available upon the acceptance of this manuscript) (Section Materials).
- We conducted comprehensive experiments to evaluate `SLR`, and provided case study on recommended drugs (Section Experimental Results). Our experiments demonstrate promising performance of `SLR`.

### Definitions and Notations

In this manuscript, we use *d* to represent a drug, and a binary matrix *A* ∈ ℝ^*m×n*^ to represent prescription data. Each row of *A*, **a**_*i*_ (*i* = 1, …, *m*), represents a prescription, and each column of *A* corresponds to a drug. Thus, If a prescription **a**_*i*_ contains a drug *d*_*j*_, the *j*-th entry in **a**_*i*_ (i.e., *a*_*i,j*_) will be 1, otherwise 0. When no ambiguity arises, the terms “a prescription” and “a binary vector **a**” are used exchangeably, and “a set of prescriptions” and “a matrix *A*” are also used exchangeably. In addition, a label *y*_*i*_ is assigned to **a**_*i*_ to indicate whether **a**_*i*_ induces a certain ADR (denoted as *y*_*i*_ = 1 or 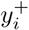) or not (denoted as *y*_*i*_ = −1 or 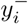).

### Joint SLIM and LogR Model: SLR

Supplementary materials present the background on `SLIM` and `LogR` model, and related work on computational methods for DDI and ADR studies. Sparse Linear Method (`SLIM`)^4^ is an efficient and state-of-the-art algorithm for top-*N* recommendation that was initially designed for e-commerce applications. In the drug recommendation problem, given a drug prescription **a**_*i*_, `SLIM` models the score of how likely an additional drug *d*_*j*_ should be co-prescribed with **a**_*i*_ as a sparse linear aggregation of the drugs in **a**_*i*_, that is,

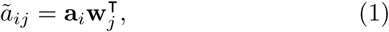

where *ã*_*ij*_ is the estimated score of *d*_*j*_ in **a**_*i*_, and 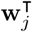 is a sparse column vector of aggregation coefficients. Note that *a*_*ij*_ = 0, that is, *d*_*j*_ is not included **a**_*i*_ originally. Drugs with high scores calculated as above will be recommended to the prescription. Thus, the scores are referred to as recommendation scores, and a prescription composed of **a**_*i*_ and a recommended drug *d*_*j*_ is referred to as a new prescription with respect to **a**_*i*_, denoted as **a**_*i*_ *∪ {d*_*j*_*}*.

We propose a new model to conduct to-avoid and safe drug recommendation to existing prescriptions. The model recommends a set of additional to-avoid drugs to an existing prescription such that each of the recommended drugs should not be taken together with the prescription in order to avoid ADRs. The model also recommends safe drugs that can be taken together with the prescription. This novel model consists of a drug recommendation component and an ADR prediction component. The recommendation component is instantiated from `SLIM` and the ADR prediction component uses logistic regression `LogR`. This model is referred to as joint sparse linear recommendation and logistic regression model and denoted as `SLR`. `SLR` learns the `SLIM` and `LogR` components jointly through solving the following optimization problem:

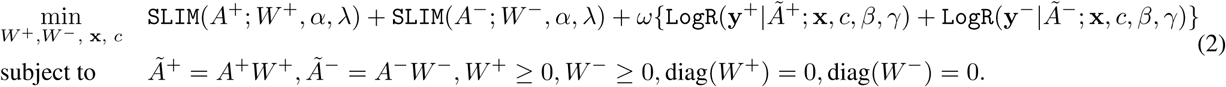

In `SLR`, *A*^+^/*A*^−^ is a set of training prescriptions known to induce ADRs/not to induce ADRs, respectively, and *Ã*^+^/*Ã*^−^ is the respective estimation from a `SLIM` model (Equation 1), that is, *Ã*^+^ = *A*^+^*W*^+^ and *Ã*^−^ = *A*^−^*W* ^−^, where *W*^+^ and *W* ^−^ are the `SLIM` parameters. The optimization algorithm for problem 2 is presented in the supplementary materials^5^.

### Learning Co-Prescription Patterns

`SLR` learns drug co-prescription patterns using its `SLIM` component (i.e., **ã** = **a***W*, or explicitly, *ã*_*i,j*_ =Σ_*k,k*≠*j*_ *a*_*i,k*_*W*_*k,j*_), that is, whether a drug should be included in a prescription is modeled as a linear function of other drugs that are included in the prescription. This modeling scheme is motivated by the existence of strong co-prescription drug pairs as we observe from real data (Section Co-Prescription Patterns on page 8). Meanwhile, linearity is a simplistic relation to model the co-prescription patterns, in which larger coefficients represent stronger co-prescription relations. Note that the zero-diagonal constraint on the coefficient matrix *W* in `SLIM` (i.e., diag(*W*^+^) = 0, diag(*W* ^−^) = 0 in Equation 2) excludes the possibility that a drug is co-prescribed with itself. The non-negativity constraint on *W* (i.e., *W*^+^ *≥* 0, *W* ^−^ *≥* 0 in Equation 2) ensures that the learned relation is on drug co-appearance. The regularization term 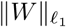 induces sparsity in *W* because not all the drugs are co-prescribed.

In `SLR`, the patterns in ADR inducing and non-ADR inducing prescriptions are modeled using two `SLIM` models (i.e., *Ã*^+^ = *A*^+^*W*^+^ and *Ã*^−^ = *A*^−^*W* ^−^) because it is expected these two patterns are different, and thus the prospectively learned *W*^+^ and *W* ^−^ will present different patterns. The pattern difference is indicated during the data pre-processing (Section Materials on page 5, Table 1). The *W* coefficients in the `SLIM` component are learned by minimizing the difference between *A* and *Ã* estimated from `SLIM`. Note that *A* is a binary matrix but *Ã* can have floating values rather than 0 and 1. Once the linear aggregation parameter *W* is learned, it can be used to recommend additional drugs that are likely to be co-prescribed with an existing prescription.

**Table 1:** Data Statistics^6^

| | $A_{\text{FAERS}}$ | | | | $A_*$ | | | |
| --- | --- | --- | --- | --- | --- | --- | --- | --- |
| | $A^-$ | | $A^+$ | | $A^-$ | | $A^+$ | |
| | $A_-^-$ | $A_0^-$ | $A_0^+$ | $A_+^+$ | $A_-^-$ | $A_0^-$ | $A_0^+$ | $A_+^+$ |
| #$\{\mathbf{a}\}$ | 621,449 | 1,264 | 8,986 | 27,387 | 2,200 | 1,264 | 2,464 | 1,000 |
| #$\{d\}$ | 1,209 | 417 | 881 | 1,201 | 562 | 417 | 692 | 679 |
| avgOrd | 6.100 | 2.351 | 3.588 | 7.096 | 2.678 | 2.351 | 3.809 | 7.615 |
| avgFrq | 1.761 | 225.317 | 13.730 | 1.402 | 42.082 | 225.317 | 20.565 | 5.520 |
| avgOR | - | 0.546 | 16.343 | - | - | 0.546 | 31.998 | - |

In this table, “# $\{\mathbf{a}\}$ ” and “# $\{d\}$ ” represent the number of prescriptions and the number of involved drugs, respectively. In each set of prescriptions, “avgOrd”/“avgFrq”/“avgOR” is the average number of drugs/average frequency/average OR of all prescriptions.

### Predicting ADR Labels

`SLR` uses its `LogR` component (i.e., *p*(*y*_*i*_ *|***ã**_*i*_; **x**, *c*) = (1 + exp(*-y*_*i*_ (**ã**_*i*_ **x^T^**+ *c*)))^-1^) to predict whether a prescription will induce ADRs or not. `LogR` produces a probability of ADR induction from a linear combination of drugs in a prescription (Equation 3). Note that the estimated prescriptions *Ã* from `SLIM` rather than *A* is used to train `LogR`. This connects `SLIM` and `LogR` to further enforce that the `SLIM` component learns co-prescription patterns that better correlate with their ADR induction labels (i.e., **y** in Equation 2). Meanwhile, the use of *Ã* also generalizes the ability of `LogR` to predict for new prescriptions, as *Ã* will have new drugs compared to *A*.

### SLR **Drug Recommendation**

`SLR` recommends drugs for each existing prescription **a** following a five-step procedure, resulting in four different recommendation methods. We will discuss these steps and the corresponding recommendation methods in detail later in this section. Note that in `SLR`, each recommended drug is considered individually together with the prescription **a**. It is also possible in real practice that multiple drugs all together should be avoided as they, together with the prescription, may induce ADRs. However, this problem is significantly more difficult as the combinatorial synergy of high-order drug combinations is highly non-trivial to estimate. This problem will be tackled in our future research.

#### Step 1: Recommendation from SLIM component

Given a prescription **a**, the recommendation scores of all drugs with respect to **a** are calculated as **ã**^+^ = **a***W*^+^ and **ã**^−^ = **a***W* ^−^, that is, **ã**^+^/**ã**^−^, the scores of being potential to-avoid/safe drugs, are calculated from *W*^+^/*W* ^−^. Based on **ã**^+^ and **ã**^−^, the top-*N* scored drugs that are not in prescription **a** will be selected as to-avoid and safe drug candidates for **a**, respectively. This set of *N* drugs is denoted as 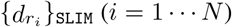. The recommendation process in this step is identical to that in the original `SLIM` method^4^. However, the `SLIM` component is learned from `SLR` (Equation 2). This recommendation method, together with the SLR learning method, is referred to as `SLR`-s.

#### Step 2: Removing low-frequency recommendations

The drugs in 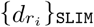 (i.e., the recommendation from the `SLIM` component in `SLR`) are removed if their co-prescription frequency with **a** is lower than a threshold *η*. Specifically, for each drug 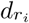 and each drug *d* ∈ **a**, we count how many times both of them are prescribed together within a prescription and sum up the counts over all the drugs in **a** as the co-prescription frequency for drug 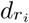. The intuition underlying the low-frequency drug removal is to explicitly follow the co-prescription patterns that have been often observed in standard of care. The reduced drug recommendation list is denoted as 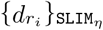.

#### Step 3: Recommendation from LogR component

For a prescription **a**, each of all possible drugs that are not in **a** is first combined with **a** to form a new prescription. Then the `LogR` component is applied on the new prescription to calculate its probability of inducing ADRs (Equation 3). The drugs that lead to the top-*N* highest/lowest predicted probabilities are recommended as the to-avoid/safe drugs. Note that the `LogR` component is learned from `SLR`. The generated recommendation list is denoted as 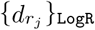.

#### Step 4: Recommendation based on drug indications

In addition to recommendations from `SLIM` and `LogR` components, we recommend another list of drugs based on drug indications. The intuition is that drugs that have similar indications tend to be prescribed together. The supplementary materials^5^ presents some examples of co-prescribed drugs with similar indications. For each drug *d* ∉ **a** and each drug *d*_*i*_ ∈ **a**, we first calculate the number of common indications between *d* and *d*_*i*_, and sum up all such numbers over *d*_*i*_ as the number of common indications for *d* with **a**. The drugs which have the top-*N* most common indications with **a** form a recommendation list, denoted as 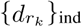. The drug indication information is extracted from the Side Effect Resource (SIDER; http://sideeffects.embl.de/).

#### Step 5: Combining recommendations

To recommend top-*N* to-avoid/safe drugs, we combine drugs from the list generated by different components, leading to the following four methods. Figure 1 demonstrates the methods.

**Figure 1:**
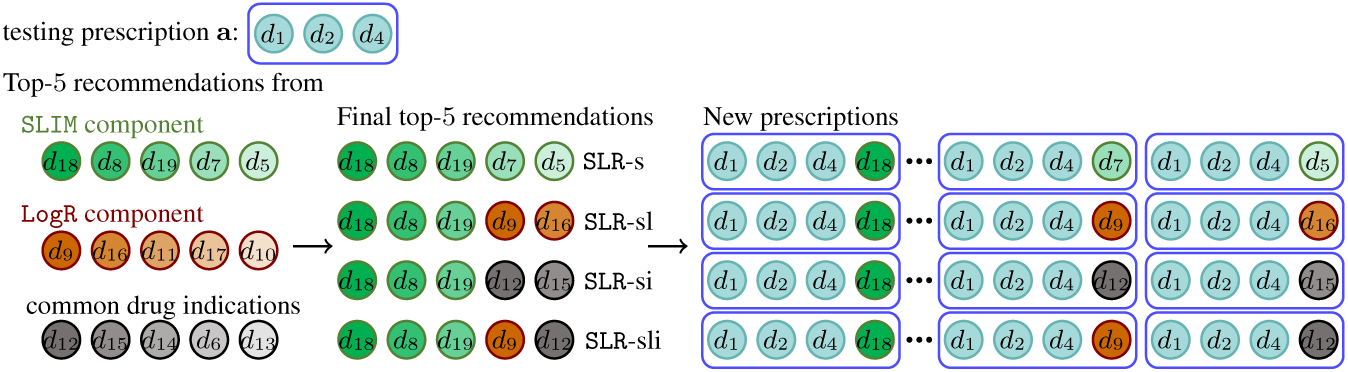
`SLR` recommendation methods

- `SLR`-s: use the recommendation list 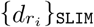 generated from Step 1 as the final recommendation list;
- `SLR`-sl: use the top-*N* drugs from the concatenation of 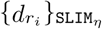 and 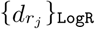 as the final recommendation list.
- `SLR`-si: use the top-*N* drugs from the concatenation of 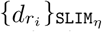 and 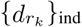 as the final recommendation list.
- `SLR`-sli: use the top-*N* drugs from the concatenation of 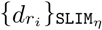, the top 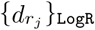 and the top 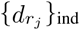 as the final recommendation list.

### Comparison Methods

#### Random Method (Rand)

In the first comparison method, we fully randomly recommend drugs for each prescription as to-avoid drugs and safe drugs, respectively. This method is referred to as random method and denoted as `Rand`. `Rand` serves as a baseline in which no learning is involved.

#### Logistic Regression Only (LogR)

In this comparison method, only a `LogR` model is learned to predict ADR labels. Prescriptions that induce ADRs are used as positive training data, and prescriptions that do not induce ADRs are used as negative training data. No `SLIM` models are learned in this method. The recommendation in this baseline method is the same as the procedure to generate a `LogR` recommendation list in `SLR` (Section Joint `SLIM` and `LogR` Model: `SLR` on page 4). This method is referred to as logistic regression only and denoted as `LogR`.

#### Sparse Linear Method Only (SLIM)

In this method, a `SLIM`^+^ model is learned on ADR inducing prescriptions, and a separate `SLIM`^−^ model is learned on non-ADR inducing prescriptions. No `LogR` is learned. For a prescription, the top drugs recommended from `SLIM`^+^ are considered as to-avoid drugs, and the top drugs recommended from `SLIM`^−^ are considered as safe drugs. This method is referred to as `SLIM` only and denoted as `SLIM`.

#### Separate Models (S+LR)

In this method, a `SLIM` and a `LogR` model are learned separately, and used together for recommendation. Specifically, a `SLIM`^+^ and a `SLIM`^−^ as in `SLIM` are learned first. Meanwhile, a `LogR` model as in `LogR` is also learned independently. The top recommendations are then generated by combining recommendations from `SLIM`^+^/`SLIM`^−^ and `LogR` model in an identical way as in `SLR`-sl. The difference is that `SLIM` and `LogR` are learned independently, not jointly as in `SLR`. This method is referred to as separate `SLIM`-`LogR` model and denoted as `S`+`LR`.

## Materials

### Training Data Generation

We use the prescription dataset *A*_FAERS_ that we generated from FDA Adverse Event Reporting System (FAERS) ^1^ in our previous research^6^ and consider myopathy as the particular ADR of interest in this work. Detailed description of dataset generation protocol is available in section 5.1 of^6^. Table 1 summarizes the dataset. Specifically, *A*_FAERS_ consists of four subsets of prescriptions: 1). 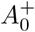 has all the prescriptions that are reported both with myopathy (case events) among some patients and without myopathy (control events) among other patients, and the Odds Ratio (`OR`) between these two cases is above 1, indicating that more possibly these prescriptions will induce myopathy; 2). 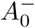 has all the prescriptions that are also reported both with and without myopathy as those in 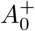, but the `OR` is below 1, indicating that more possibly these prescriptions will not induce myopathy; 3). 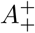 consists of prescriptions that only have reports of myopathy (i.e., only in case events); and 4). 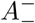 consists of prescriptions that only have reports of non-myopathy (i.e., only in control events). The set of prescriptions in case events is denoted as *A*^+^ (i.e., 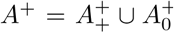) and is used as the positive set in *A*_FAERS_. The set of prescriptions in control events is denoted as *A*^−^ (i.e., 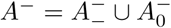) and is used as the negative set in *A*_FAERS_.

To use more frequent and more confident prescriptions for model training, we further generated a prescription dataset from *A*_FAERS_. The detailed generation protocol is presented in section 5.2 in^6^. This dataset is denoted as *A*_*\**_ and summarized in Table 1. Note that *A*_*\**_ is the set of labeled prescriptions that are used for model training.

### Evaluation Protocols and Metrics

#### Five-Fold Cross Validation

The performance of different methods is evaluated via five-fold cross validation. The dataset *A*_*\**_ is randomly split into five folds of equal size. Four folds are used for model training and the rest fold is used for testing. The corresponding training set is denoted as 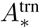. In the fold for testing, each prescription of order *k* (*k* = 2, 3, *…*) is first duplicated into *k* copies. Then each of the *k* drugs is excluded from each of the *k* duplicates, respectively, to generate a testing prescription of order *k*-1. We further unify these testing prescriptions and exclude those testing prescriptions that appear in 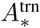. The remaining testing prescriptions constitute the testing set, denoted as 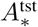. Figure 2 demonstrates the testing set generation process. The drug recommendation will be conducted and evaluated on 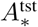 prescriptions for five times, with one fold for testing each time. The final result is the average of the five experiments.

**Figure 2:**
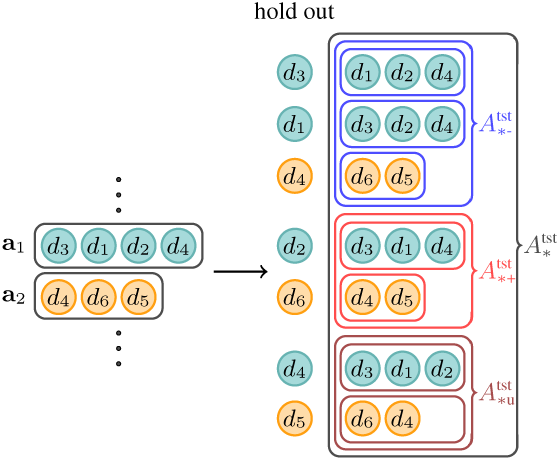
Testing set generation

#### Knowledge Pool Generation

To evaluate the performance of various methods, we first define a knowledge pool of labeled prescriptions, denoted as *A*_pool_, which the recommended new prescriptions can be searched from and evaluated against. This knowledge pool serves as the entire knowledge space which is unknown during training. For a training set 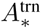, we consider 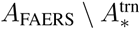 as the knowledge pool *A*_pool_. 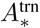 is excluded from *A*_pool_ because we can always search testing prescriptions in 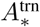, where prescriptions have known labels at the time of training, in order to identify corresponding to-avoid prescriptions in 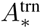, and therefore no recommendations will be needed. By excluding 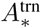 from *A*_pool_, we care about recommending particularly to-avoid/safe drugs to prescriptions such that the new prescriptions have not been observed before. We believe this application is interesting and significant in real practice.

#### Evaluation Metrics

Each of the methods will generate *N* recommended to-avoid/safe drugs for each testing prescription. Given a knowledge pool *A*_pool_, the performance of these methods is evaluated using *Normalized Recall on Positives/Negatives*, denoted as 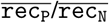, defined as follows,

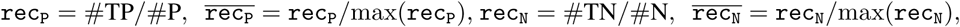

where #TP/#TN is the number of recommended to-avoid/safe prescriptions for 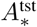 that have positive/negative labels in *A*_pool_, #P/#N is the number of all possible ground-truth to-avoid/safe prescriptions for 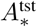 that have positive/negative labels in *A*_pool_, and max(`rec`_P_)/max(`rec`_N_) is the maximum possible `rec`_P_/`rec`_N_ value. In particular, max(rec_P_)/max(rec_N_) is achieved when all the top-*N* recommendations lead to true-positive/true-negative (to-avoid/safe) prescriptions. Note that max(rec_P_)/max(rec_N_) can be smaller than 1. This is because for each testing prescription, the number of recommended prescriptions is limited to *N* and can be smaller than the number of all its possible ground-truth to-avoid/safe prescriptions in *A*_pool_. Thus, 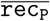 and 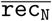 measure how much each method can achieve to its best. Higher 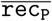 and 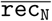 values indicate better performance. In particular, higher 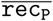 indicates stronger performance on recommending to-avoid drugs, and higher 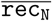 indicates stronger performance on recommending safe drugs. The maximum possible values of rec_P_ and rec_N_ (i.e., max(rec_P_) and max(rec_N_)) are presented in Table S2 in the supplementary materials.

The third metric is the harmonic mean of 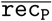 and 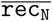, that is, 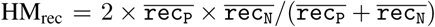, and is denoted as HM_rec_. Higher HM_rec_ values indicate better performance. Note that we did not use conventional ranking-based metrics such as average precision at *k* because we believe in clinical practice of multi-drug prescription, people care whether all the to-avoid drugs can be correctly recommended (i.e., a notion of recall) much more than how the to-avoid drugs are ranked.

## Experimental Results

### Overall Comparison

We present the overall performance comparison of all the methods in Table 2. Note that Table 2 presents the performance of different methods in terms of their best 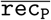, best 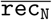 and HM_rec_, respectively, and lists the corresponding values on the other two evaluation metrics when each respective best performance is achieved. Optimal parameters are not presented in the tables due to space limit. Overall, recommendation-based methods (i.e., `SLIM`, `S`+`LR` and `SLR`-based methods) substantially outperform `LogR` and `Rand`. `LogR` learns and predicts the labels of prescriptions, but it does not learn or leverage the patterns among the drugs in prescriptions, even though `LogR` learns the labels substantially better than random (`Rand`). In terms of best 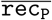, `SLIM`, `S`+`LR`, `SLR`-s, `SLR`-sl, `SLR`-si and `SLR`-sli are very comparable (all around 0.26), with `S`+`LR` 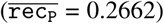 slightly better than the others. In terms of best 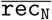, `SLR`-sl 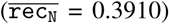, `SLR`-si and `SLR`-sli 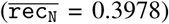 are comparable. `SLR`-si and `SLR`-sli are 5.3% better than `S`+`LR` and 6.2% better than `SLIM`, where both of `S`+`LR` and `SLIM` learn a `SLIM` component independently. In terms of best HM_rec_, `SLR`-sl (HM_rec_ =0.3127) and `SLR`-sli (HM_rec_ = 0.3136) are also comparable. `SLR`-sli is 2.0% better than `S`+`LR` and 3.1% better than `SLIM`. Overall, `SLR`-sl and `SLR`-sli are very comparable, and both are slightly better than `SLR`-s and `SLR`-si. `SLR`-based methods (i.e., `SLR`-s, `SLR`-sl, `SLR`-si and `SLR`-sli) outperform `SLIM`, a strong baseline for top-*N* recommendation but learned without consideration of recommendation labels. `SLR`-based methods also outperform `S`+`LR`, which considers recommendation labels but learns labels independently of recommendations. These results demonstrate the strong potential of `SLR` methods in recommending to-avoid/safe drugs for existing prescriptions. Detailed parameter study is presented in Section Parameter Study on page 3 and Section Top-*N* Performance on page 4 in the supplementary materials^5^.

**Table 2:** Overall Performance Comparison on 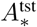

| mdl | best $\overline{\text{rec}}_P$ | | | best $\overline{\text{rec}}_N$ | | | best $\text{HM}_{\text{rec}}$ | | |
| --- | --- | --- | --- | --- | --- | --- | --- | --- | --- |
| | $\overline{\text{rec}}_P$ | $\overline{\text{rec}}_N$ | $\text{HM}_{\text{rec}}$ | $\overline{\text{rec}}_P$ | $\overline{\text{rec}}_N$ | $\text{HM}_{\text{rec}}$ | $\overline{\text{rec}}_P$ | $\overline{\text{rec}}_N$ | $\text{HM}_{\text{rec}}$ |
| Rand | 0.0020 | 0.0003 | 0.0006 | 0.0020 | 0.0003 | 0.0006 | 0.0020 | 0.0003 | 0.0006 |
| LogR | 0.1621 | 0.2320 | 0.1908 | 0.1614 | 0.2397 | 0.1927 | 0.1614 | 0.2397 | 0.1927 |
| SLIM | 0.2564 | 0.3739 | 0.3041 | 0.2072 | 0.3747 | 0.2668 | 0.2564 | 0.3739 | 0.3041 |
| S+LR | **0.2662** | 0.3604 | 0.3061 | 0.2574 | 0.3777 | 0.3061 | 0.2649 | 0.3666 | 0.3074 |
| SLR-s | 0.2576 | 0.3739 | 0.3049 | 0.2468 | 0.3823 | 0.2999 | 0.2574 | 0.3755 | 0.3053 |
| SLR-sl | 0.2633 | 0.3381 | 0.2959 | 0.2608 | 0.3910 | 0.3127 | 0.2608 | 0.3910 | 0.3127 |
| SLR-si | 0.2583 | 0.3943 | 0.3121 | 0.2472 | **0.3978** | 0.3048 | 0.2583 | 0.3943 | 0.3121 |
| SLR-sli | 0.2654 | 0.3836 | 0.3136 | 0.2474 | **0.3978** | 0.3050 | 0.2654 | 0.3836 | **0.3136** |

The column "mdl" corresponds to models. The best overall performance is **bold**.

**Table 3:** Performance Comparison on 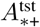

| mdl | best $\overline{\text{rec}}_P$ | | | best $\overline{\text{rec}}_N$ | | | best $\text{HM}_{\text{rec}}$ | | |
| --- | --- | --- | --- | --- | --- | --- | --- | --- | --- |
| | $\overline{\text{rec}}_P$ | $\overline{\text{rec}}_N$ | $\text{HM}_{\text{rec}}$ | $\overline{\text{rec}}_P$ | $\overline{\text{rec}}_N$ | $\text{HM}_{\text{rec}}$ | $\overline{\text{rec}}_P$ | $\overline{\text{rec}}_N$ | $\text{HM}_{\text{rec}}$ |
| Rand | 0.0010 | 0.0003 | 0.0004 | 0.0010 | 0.0003 | 0.0004 | 0.0010 | 0.0003 | 0.0004 |
| LogR | 0.1964 | 0.1796 | 0.1875 | 0.1964 | 0.1796 | 0.1875 | 0.1964 | 0.1796 | 0.1875 |
| SLIM | 0.2519 | 0.4110 | 0.3123 | 0.2519 | 0.4110 | 0.3123 | 0.2519 | 0.4110 | 0.3123 |
| S+LR | 0.2528 | 0.4046 | 0.3111 | 0.2528 | 0.4046 | 0.3111 | 0.2528 | 0.4046 | 0.3111 |
| SLR-s | **0.2530** | 0.4111 | 0.3131 | 0.2486 | 0.4138 | 0.3104 | 0.2530 | 0.4111 | 0.3131 |
| SLR-sl | 0.2529 | 0.4119 | 0.3133 | 0.2493 | 0.4113 | 0.3104 | 0.2529 | 0.4119 | 0.3133 |
| SLR-si | 0.2528 | 0.4150 | 0.3141 | 0.2482 | **0.4179** | 0.3112 | 0.2528 | 0.4150 | **0.3141** |
| SLR-sli | **0.2530** | 0.4131 | 0.3137 | 0.2482 | **0.4179** | 0.3112 | 0.2530 | 0.4131 | 0.3137 |

The column "mdl" corresponds to models. The best overall performance is **bold**.

### Comparison on Different Testing Subsets

Among all the testing prescriptions in 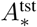, some prescriptions can be found in *A*_FAERS_ along with known ADR labels, while the others have unknown ADR labels. We separate 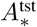 into the following subsets according to their labels in *A*_FAERS_: (1). 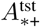: the set of prescriptions in 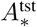 that have positive labels in *A*_FAERS_; (2). 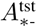: the set of prescriptions in 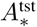 that have negative labels in *A*_FAERS_; (3). 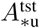: the set of prescriptions in 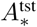 that do not have labels in *A*_FAERS_. Thus, 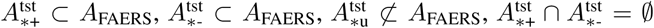 and 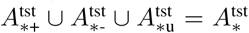. Note that if a prescription with an unknown label contains only one drug and can be found in 𝒟_Myo_ (i.e., the set of drugs which induce myopathy on their own), we still categorize it as in 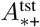. Different subsets here correspond to different concerns of interest. For 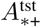, we are more concerned with safe drug recommendations (i.e., 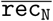 in evaluation) so as not to further increase the ADR risks of prescriptions. For 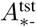, we focus more on to-avoid drug recommendation (i.e., 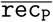 in evaluation) so as not to introduce ADRs in prospective prescriptions. For 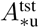, we are interested in both to-avoid and safe drug recommendations. Table 4 presents the statistics of different sets of testing prescriptions. In particular, the column 𝒟_Myo_(%) in Table 4 presents the average percentage of the 𝒟 _Myo_ drugs in a prescription. In specific, we first calculate the percentage of 𝒟 _Myo_ drugs in each prescription and take the average. 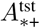, in which all the prescriptions are positive, contains the most 𝒟 _Myo_ drugs on average.

**Table 4:**
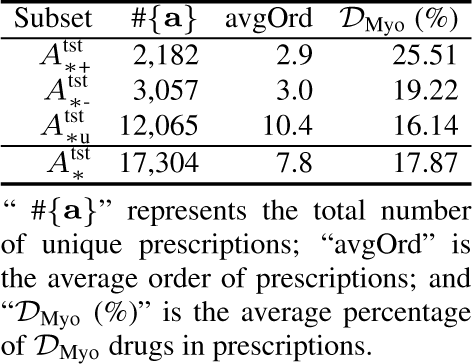
Statistics on Different Testing Subsets

#### Performance Comparison on 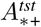

Table 3 presents the performance of different methods on 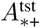. In terms of best 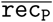, `SLR`-based methods and `S`+`LR` have similar performance (around 0.2528) and slightly better than `SLIM`. In terms of best 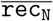, `SLR`-based methods and `SLIM` have similar performance (around 0.4100), with SLR-si and `SLR`-sli slightly better than the others. In terms of best HM_rec_, the `SLR`-based methods are similar to `SLIM`. These results show that for 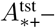 the set of prescriptions that induce ADRs, joint learning `SLIM` and `LogR` does not introduce additional benefits on top of `SLIM`, although all these methods are significantly better than `Rand` and `LogR`. However, comparing the best 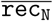 values in Table 3 and in Table 2, `SLR`-based methods have better performance (best 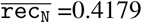) on 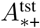 than on 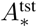 (best 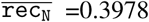). This indicates that for ADR-inducing prescriptions, our methods are able to recommend safe drugs that taken together with the ADR-inducing prescriptions would actually lead to no/less ADR effects.

#### Performance Comparison on 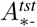

Table 5 presents the performance of different methods on 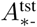. The `SLR`-based methods and `SLIM` have a similar performance in their best 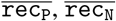 and HM_rec_. However, comparing the results in Table 5 with the results in Table 3 and Table 2, `SLR`-based methods can achieve even better 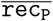 values 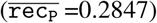 on 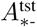 than those on 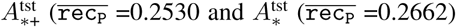. As for 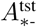, we care more on recommending to-avoid drugs. The better 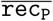 values on 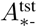 indicate the capability of `SLR`-based methods in recommending to-avoid drugs that taken together with the non ADR-inducing prescriptions would not induce ADRs. `SLR`-based methods also have better performance on 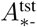 in terms of 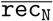 and HM_rec_, compared to that on 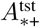 and 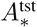.

**Table 5:**
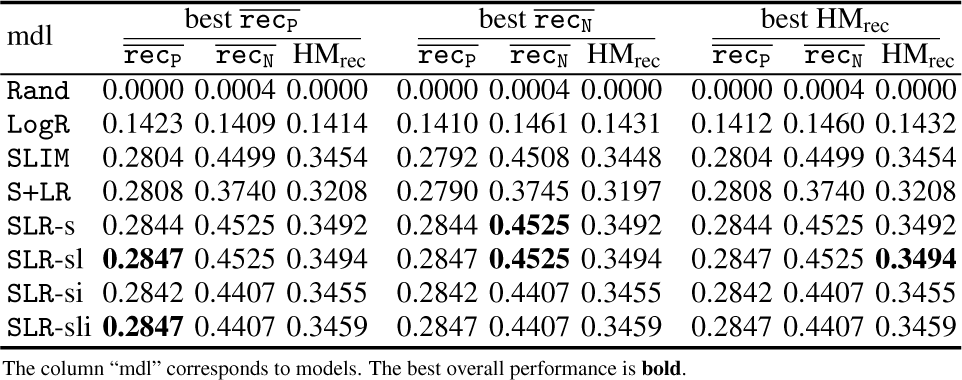
Performance Comparison on 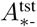

#### Performance Comparison on 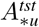

Table 6 presents the performance of different methods on 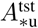. Different from the performance on 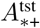 and 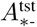, on 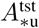, `S+LR` has the best performance in terms of 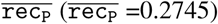 compared to `SLR`-sli, the best performing `SLR`-based methods 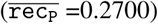. `SLIM` is worse than `SLR`-based methods in terms of 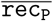. However, in terms of best 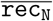, on 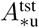, `SLR`-sli and `SLR`-sl are significantly better than other methods, followed by `SLR`-si and `SLR`-s, while `SLIM` and `LogR` do not perform comparably. A similar trend holds in terms of best HM_rec_. The difference of the performance trends on 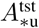 may be due to the different characteristics of 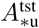 (Table 4). For example, the larger size of drug combinations may allow `LogR` alone to better predict prescription labels, particularly the positive labels. Overall, on 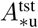, `SLR`-based methods outperform `SLIM` and `LogR` on average. This indicates the capability of `SLR`-based methods in recommending safe drugs when the testing prescriptions do not have ADR labels.

**Table 6:**
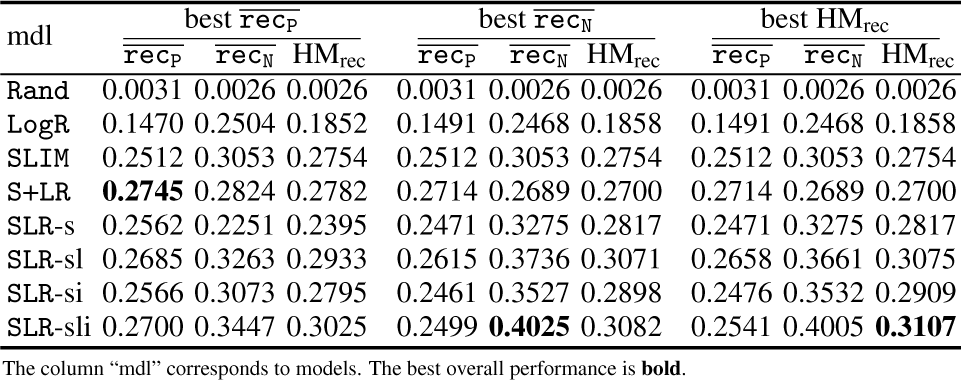
Performance Comparison on 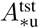

### Recommendation Method Comparison

Table 7 presents the contribution from various recommendation components of `SLR`-sli (i.e., the `SLIM` component, the `LogR` component and the indication component as in Figure 1). In Table 7, the average number of true positive prescriptions that are recommended by `SLR`-sli for 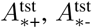 and 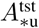 in terms of best HM_rec_ is 503.0, 249.0, and 901.8, respectively, and the corresponding average number of true negative prescriptions are 1204.0, 1234.2, and 829.8, respectively. For 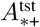 and 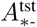, the recommended true positives are fewer than the true negatives. This may be due to the fact that, as in Table 1, the knowledge pool *A*_FAERS_ has more negative prescriptions, which leads to a higher probability of a recommended prescription being true negative in *A*_FAERS_. However, for 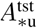, there are more recommended true positives than true negatives. The 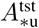 testing prescriptions have higher average order than the 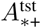 and 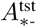 testing prescriptions as in Table 4. Such higher order leads to higher risks of those 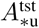 testing prescriptions on their own being true positives (as in Table 1, true positives tend to have higher order than true negatives). The fact that `SLR`-sli recommends more true positives than true negatives for 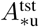 demonstrates its strong capability of identifying to-avoid drugs particularly when the existing prescriptions already have high risks of ADRs.

**Table 7:**
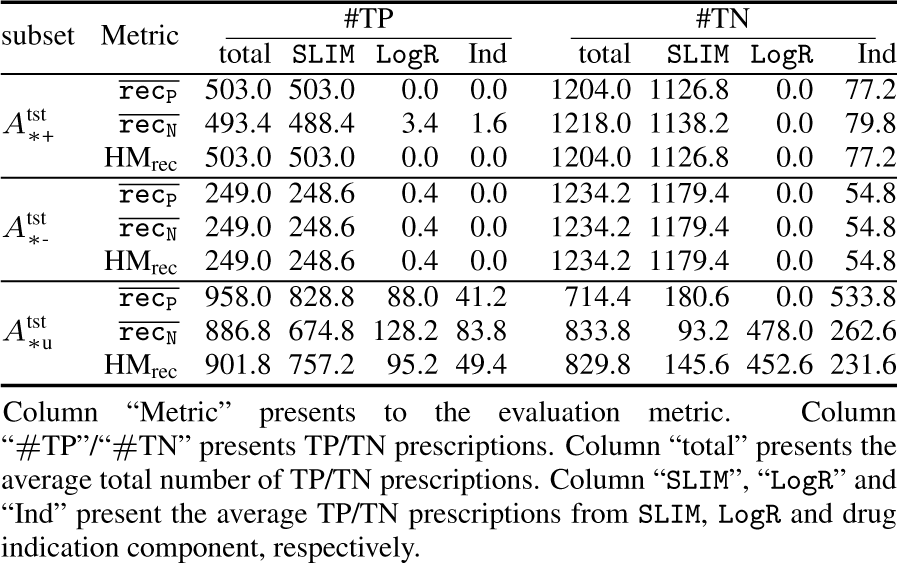
Recommendation Statistics from `SLR`-sli

For 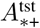 and 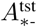, almost all the true positive and true negative recommendations are from the `SLIM` component of `SLR`-sli. However, for 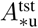, the recommendations are contributed from all of the three components (e.g., 757.2, 95.2, and 49.4 true positives from `SLIM`, `LogR` and indication component, respectively). This indicates that for high-order prescriptions, to-avoid/safe drug recommendation may require the collective consideration of various factors including the drug coprescription patterns and drug indications, etc. This also demonstrates the capability of `SLR`-sli in leveraging the recommendation lists strategically from various recommendation components in order to generate reliable to-avoid/safe drug recommendations.

### ADR-Relevant Drug Identification

We analyze how `SLR`-sli can identify ADR-relevant single drugs using **x** in the `LogR` component. The **x** values represent how strong the association between the corresponding drug and the ADR label is. The larger positive/negative value of *x*_*i*_ indicates higher possibility for drug *d*_*i*_ in inducing/not inducing ADR. The top-5 largest positive and negative weights in the **x** from `SLR`-sli and the corresponding drugs are presented in Table 8. Note the largest positive/negative weights correspond to drugs that are most predictive of ADR/non-ADR induction, and the `SLR`-sli corresponds to the best HM_rec_ performance on 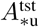 as in Table 7, where `LogR` contributes most. In Table 8, it turns out only two **x** values are positive, corresponding to drug atorvastatin and simvastatin. These two drugs have been reported in SIDER as to induce myopathy, and 28.72% of the ADR inducing prescriptions have at least one of these two drugs. This indicates that the `LogR` component in `SLR`-sli is able to identify single drugs that are most ADR relavant. Meanwhile, the top-5 largest negative weights correspond to warfarin, ibuprofen, metformin, quetiapine, and valproic acid. In *A*_*\**_, the frequency of these 5 drugs in *A*^+^ is 119, 110, 176, 46, and 54, respectively, and much higher than the average frequency 14.05 of all the drugs; the frequency of these 5 drugs in *A*^−^ is 124, 86, 148, 102, and 95, respectively, and also much higher than the average frequency 7.32 of all the drugs. This indicates that the `LogR` component in `SLR`-sli is also able to identify single drugs that are most ADR safe.

**Table 8:** Most Indicating Drugs from `SLR`-sli

| Top-5 |  | Bottom-5 |  |
| --- | --- | --- | --- |
| $x$ | $d$ | $x$ | $d$ |
| 7.944 | **atorvastatin** | -10.783 | warfarin |
| 6.430 | **simvastatin** | -9.492 | ibuprofen |
| - | - | -3.592 | metformin |
| - | - | -1.761 | quetiapine |
| - | - | -0.673 | valproic acid |

“ $x$ ” is the entry value of drug in $x$ ; “ $d$ ” corresponds to the drug; and “Top-5” and “Bottom-5” correspond to the top-5 largest and smallest values in $x$ , respectively. Drugs in SIDER to induce myopathy on their own are **bold**.

### Co-Prescription Patterns

We analyze the co-prescription patterns using *W*^+^ and *W* ^−^ from `SLR`-sli of the best HM_rec_ performance. *W*^+^/*W* ^−^ captures the co-prescription patterns among drugs that often appear in ADR inducing/no-ADR inducing prescriptions, respectively. The values in *W*^+^/*W* ^−^ can be considered as measurement of such patterns: higher/lower values correspond to stronger/weaker co-prescription patterns. The top-10 largest values in *W*^+^ and *W* ^−^ from the optimal `SLR`-sli model and the corresponding co-prescribed drug pairs on 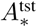 are presented in Table 9.

**Table 9:** Co-Prescription Patterns from `SLR`-sli on 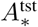

| $W^+$ | | | | $W^-$ | | | |
| --- | --- | --- | --- | --- | --- | --- | --- |
| $w$ | $d_1$ | $d_2$ | freq | $w$ | $d_1$ | $d_2$ | freq |
| 0.7021 | ethinyl estradiol | etonogestrel | 38 | 0.6275 | ethinyl estradiol | norelgestromin | 23 |
| 0.5903 | hydrochlorothiazide | triamterene | 32 | 0.5932 | **emtricitabine** | **tenofovir** | 35 |
| 0.5874 | sulfamethoxazole | trimethoprim | 31 | 0.5916 | acetaminophen | hydrocodone | 36 |
| 0.5719 | acetaminophen | hydrocodone | 133 | 0.5523 | fluticasone propionate | salmeterol | 35 |
| 0.5302 | fluticasone propionate | salmeterol | 44 | 0.5189 | lopinavir | **ritonavir** | 36 |
| 0.5034 | amoxicillin | clavulanate | 22 | 0.4668 | budesonide | formoterol | 17 |
| 0.4967 | carbidopa | levodopa | 17 | 0.4496 | fluorouracil | leucovorin | 18 |
| 0.4595 | acetaminophen | codeine | 27 | 0.4459 | conjugated estrogens | medroxyprogesterone | 17 |
| 0.4492 | **emtricitabine** | **tenofovir** | 22 | 0.4104 | buprenorphine | naloxone | 9 |
| 0.4268 | pamidronate | zoledronate | 42 | 0.3837 | amphetamine | dextroamphetamine | 8 |

In this table, “ $w$ ” is the value of the entry corresponding to the drug pairs in $W^+/W^-$ ; “ $d_1$ ” and “ $d_2$ ” are the two drugs in the drug pair, and “freq” is the frequency of the corresponding drug pairs in training data $A^+$ and $A^-$ . Drugs that are reported in SIDER to induce myopathy on their own are **bold**.

In Table 9, the learned *W*^+^ has a density 10.89%, and its top-5 largest values correspond to the following drug pairs: (ethinyl estradiol, etonogestrel), (hydrochlorothiazide, triamterene), (sulfamethoxazole, trimethoprim), (acetaminophen, hydrocodone), and (fluticasone propionate, salmeterol). These drug pairs are frequently prescribed together in *A*^+^. The number of prescriptions in *A*^+^ that contain these drug pairs is 38, 32, 31, 133, and 44, respectively (“freq” in Table 9), compared to the average number of *A*^+^ prescirptions of all possible drug pairs, which is 2.48. This demonstrates that `SLR`-sli is able to capture the co-prescription patterns among perscriptions. In addition, many of the co-prescribed drugs share similar medical purposes. For example, ethinyl estradiol and etonogestrel are often used for birth control, and acetaminophen and hydrocodone are often prescribed together to relieve moderate to severe pains and treat fever.

In Table 9, the learned *W* ^−^ has a density 18.20%, and its top-5 largest values correspond to the following drug pairs: (ethinyl estradiol, norelgestromin), (emtricitabine, tenofovir), (acetaminophen, hydrocodone), (fluticasone propionate, salmeterol), and (lopinavir, ritonavir). These drug pairs are also frequently co-prescribed in *A*^−^. The number of *A*^−^ prescriptions that have these drugs pairs is 23, 35, 36, 35 and 36, respectively, compared to 2.07, the average number of *A*^−^ prescriptions that have all possible drug pairs. These drugs pairs also share similar medical purposes. For example, emtricitabine and tenofovir, and lopinavir and ritonavir are commonly used in HIV cocktail therapy, and fluticasone propionate and salmeterol are prescribed together for preventing difficulty in breathing.

It is also noted that in Table 9, the learned *W*^+^ and *W* ^−^ values are not necessarily in order with the drug pair frequencies. This indicates that the `SLR`-sli is able to discover patterns related to ADRs that are not simply represented by prescription frequencies. In addition, the two lists of top-10 drug pairs corresponding to *W*^+^ and *W* ^−^ values have 3 common pairs (e.g., (acetaminophen, hydrocodone), (fluticasone propionate, salmeterol) and (emtricitabine, tenofovir)) and 7 different pairs, respectively. This may illustrate the common practices in both ADR-inducing and non-ADR-inducing drug perscribing, but meanwhile the different distributions of ADR-inducing and non-ADR inducting prescriptions. Co-prescription patterns corresponding to the best HM_rec_ on 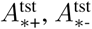 and 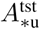 are presented in Table S6, S7, and S8, respectively, in the supplementary materials^5^.

### Case Study

Table S9 in the supplementary materials^5^ presents some examples of testing prescriptions and their recommended to-avoid drugs from `SLR`-sli such that the corresponding new prescriptions (i.e., testing prescriptions and recommended drugs together) are ADR-inducing. When the testing prescriptions already induce ADRs (i.e., in 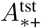; the first row block in Table S9), `SLR`-sli is able to identify additional single drugs such that when the recommended drug is taken together with the testing prescription, the `OR` of the new prescription increases over that of the testing prescription. For example, the testing prescription (fusidic acid, simvastatin) in 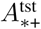 is ADR inducing with `OR` =39.809 (p-value=1.281 × 10^-14^). `SLR`-sli recommends ramipril such that (fusidic acid, simvastatin, ramipril) also induces ADR with an increased `OR` =159.2182 (p-value=1.8109 × 10^-9^). Such examples also indicate that the increased `OR` is due to the additional recommended drugs, and the potential directional interactions between the recommended drugs and the testing prescriptions. This conclusion may hint on the research on directional drug-drug interactions^7^.

When the testing prescriptions do not induce ADRs (i.e., in 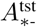; the second row block in Table S9), `SLR`-sli is able to recommend to-avoid drugs that will introduce ADRs to the corresponding new prescriptions. In particular, there are examples in which the recommended to-avoid drugs themselves are not ADR-inducing. For example, the testing prescription (dicyclomine, gabapentin, lansoprazole, lorazepam, pamidronate, zoledronate, zolpidem) is ADR negative and the recommended to-avoid drug “oxycodone” is itself ADR negative as well. However, the corresponding new prescription is ADR inducing (`OR` =34.116, p-value=7.007 × 10^-4^). Another example is for the testing prescription (acetylsalicylic acid, amlodipine, atorvastatin, isosorbide mononitrate, lisinopril, olmesartan, omeprazole, quinine). Although atorvastatin is ADR-positive, this prescription its own is ADR-negative. However, `SLR`-sli recommends another ADR-negative drug “atenolol” that results in an ADR-positive prescription (acetylsalicylic acid, amlodipine, atorvastatin, isosorbide mononitrate, lisinopril, olmesartan, omeprazole, quinine, atenolol) (`OR` =13.646, p-value=3.566 × 10^-3^). This demonstrates that the recommendations from `SLR`-sli are not trivial and do capture the interaction patterns among multiple drugs in a prescription. The same conclusion can also be derived from the examples for testing prescriptions in 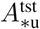 (i.e., the third row block in Table S9). For example, for the testing prescription (acetylsalicylic acid, amlodipine, esomeprazole, hydrochlorothiazide, potassium chloride, sorafenib, triamterene) in 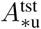, `SLR`-sli recommends “furosemide” which, taken together with the testing prescription, will induce ADR, although furosemide itself is ADR-negative. More examples on safe drug recommendation are available in Table S10 in the supplementary materials^5^.

## Discussions and Conclusions

In this manuscript, we formally formulated the to-avoid drug recommendation problem and the safe drug recommendation problem. We developed a joint sparse linear recommendation and logistic regression model (`SLR`) to tackle the recommendation problems, and developed real datasets and new evaluation protocols to evaluate the model. To the best of our knowledge, this is the first time such problems are formally formulated and corresponding computational solutions are proposed. Our experiments demonstrate strong performance of `SLR` compared to other methods. In particular, `SLR` is able to 1). recommend safe drugs that taken together with the ADR-inducing prescriptions would lead to no/less ADR effects; 2). to recommend to-avoid drugs that taken together with the non ADR-inducing prescriptions would not induce ADRs; 3). to recommend safe drugs when the testing prescriptions do not have an ADR labels; and 4). to leverage the recommendation lists strategically from various recommendation components in order to generate reliable to-avoid/safe drug recommendations. In `SLR`, we use simplistic linear models. Non-linear models (for both recommendation and label prediction) together may be able to better discover co-prescription patterns and predict labels jointly. We will investigate non-linear models in the future work. In addition, `SLR` is learned on a population level without personalization to each individual patient, because the ADR labels are generated over a population. With patient data available, we may extend the methods to personalized drug recommendation.

## Acknowledgment

This material is based upon work supported by the National Science Foundation under Grant Number IIS-1855501 and IIS-1827472.

## Footnotes

1 https://www.fda.gov/drugs/informationondrugs/ucm135151.htm

