## supplemental files for "Computational Drug Recommendation Approaches toward Safe Polypharmacy"

### Related Work

**Computational Methods for DDI and ADR Studies** Significant research efforts have focused on detecting DDIs, and can be broadly classified into four categories. Methods of the first category analyze medical literature and/or electronic medical records, and extract mentioned drug pairs[1]. Methods in the second category integrate various biochemical and molecular drug/target data to measure drug-drug similarities and score/predict pairwise DDIs. These data include phenotypic and genomic information[2], and drug side effects[3], etc. Methods of the third category leverage healthcare information on social media and online communities to detect DDIs[4]. The fourth category focuses on using numerical models to predict the dose responses to multiple drugs[5]. A recent thread is dedicated to understanding the interaction patterns among high-order DDIs, and how such patterns can relate to induced ADRs[6].

**Recommender Systems** Top-$N$ recommender systems, which recommend the top-$N$ items that are most likely to be preferred by users, have been used in a variety of applications in e-commerce and social networking, etc[7]. Recommender systems have been recently applied to prioritizing healthcare information, due to a tremendous need for personalized healthcare[8]. Current applications along this line include recommending physicians to patients on specific diseases[9], and recommending drugs for patient symptoms[10], etc. However, to the best of our knowledge, very little research has been done on new prescription recommendation particularly with ADR concerns.

### Background

**Sparse Linear Method for top-$N$ Recommendation** Sparse Linear Method (`SLIM`)[11] is an efficient and state-of-the-art algorithm for top-$N$ recommendation that was initially designed for e-commerce applications. In the drug recommendation problem, given a drug prescription $\mathbf{a}_i$, `SLIM` models the score of how likely an additional drug $d_j$ should be co-prescribed with $\mathbf{a}_i$ as a sparse linear aggregation of the drugs in $\mathbf{a}_i$, that is,

$$\tilde{a}_{ij} = \mathbf{a}_i \mathbf{w}_j^{\mathsf{T}}, \tag{1}$$

where $\tilde{a}_{ij}$ is the estimated score of $d_j$ in $\mathbf{a}_i$, and $\mathbf{w}_j^{\mathsf{T}}$ is a sparse column vector of aggregation coefficients. Note that $a_{ij} = 0$, that is, $d_j$ is not included $\mathbf{a}_i$ originally. Drugs with high scores calculated as above will be recommended to the prescription. Thus, the scores are referred to as recommendation scores, and a prescription composed of $\mathbf{a}_i$ and a recommended drug $d_j$ is referred to as a new prescription with respect to $\mathbf{a}_i$, denoted as $\mathbf{a}_i \cup \{d_j\}$. The intuition of using `SLIM` for drug recommendation will be discussed later in Section Joint `SLIM` and `LogR` Model: `SLR` on page 3.

To learn $W = [\mathbf{w}_1^{\mathsf{T}}, \mathbf{w}_2^{\mathsf{T}}, \cdots, \mathbf{w}_n^{\mathsf{T}}]$, `SLIM` solves the following optimization problem,

$$\min_{W} \quad \texttt{SLIM}(A; W, \alpha, \lambda) = \frac{1}{2}\|A - AW\|_F^2 + \frac{\alpha}{2}\|W\|_F^2 + \lambda\|W\|_{\ell_1}, \quad \text{subject to} \quad W \geq 0, \text{diag}(W) = 0, \tag{2}$$

where $\|W\|_{\ell_1} = \sum_{i=1}^{n}\sum_{i=1}^{n}|w_{ij}|$ is the entry-wise $\ell_1$-norm of $W$, and $\|\cdot\|_F$ is the matrix Frobenius norm. In `SLIM`, $W$ converts a binary $A$ into its estimation $\tilde{A}$ of floating values, which could recover unseen non-zero entries in $A$.

**Logistic Regression for Label Prediction** We can formulate the problem of predicting whether a prescription of multiple drugs induces a particular ADR as a binary classification problem, and solve the problem using logistic regression (`LogR`):

$$p(y_i|\mathbf{a}_i; \mathbf{x}, c) = (1 + \exp(-y_i(\mathbf{a}_i\mathbf{x}^{\mathsf{T}} + c)))^{-1}, \tag{3}$$

where $\mathbf{x}^{\mathsf{T}}$ and $c$ are the parameters. To learn the parameters, `LogR` solves the following optimization problem,

$$\min_{\mathbf{x},c} \quad \texttt{LogR}(\mathbf{y}|A; \mathbf{x}, c, \beta, \gamma) = \sum_{i=1}^{m}\log\{1 + \exp[-y_i(\mathbf{a}_i\mathbf{x}^{\mathsf{T}} + c)]\} + \frac{\beta}{2}\|\mathbf{x}\|_2^2 + \gamma\|\mathbf{x}\|_1, \tag{4}$$

where $\mathbf{y}$=$[y_1; y_2; \cdots; y_m]$, $\|\mathbf{x}\|_1$=$\sum_{i=1}^{n}|x_i|$, and $\|\mathbf{x}\|_2^2$=$\sum_{i=1}^{n}{x_i}^2$.

**Training** `SLR`

The optimization problem 2 can be solved through the alternating direction method of multipliers (ADMM)[12]. We introduce a new variable $Z$ and thus the following augmented Lagrangian as the new objective to optimize:

$$
\begin{aligned}
\min_{\substack{\boldsymbol{\Theta}=\{W^+,W^-,\\ Z^+,Z^-,\mathbf{x},\,c\}}} \quad & L(A^+, A^-, y^+, y^-; \boldsymbol{\Theta},\ \mathbf{u}^+,\ \mathbf{u}^-, \rho^+, \rho^-) = \\
& \texttt{SLIM}(A^+; W^+, \alpha, \lambda) + \texttt{SLIM}(A^-; W^-, \alpha, \lambda) \\
& \omega\{\texttt{LogR}(\mathbf{y}^+|\tilde{B}^+; \mathbf{x}, c, \beta, \gamma) + \texttt{LogR}(\mathbf{y}^-|\tilde{B}^-; \mathbf{x}, c, \beta, \gamma)\} \\
& \mathbf{u}^{+\mathsf{T}}\mathbf{v}^+ + \frac{\rho^+}{2}\|\mathbf{v}^+\|_2^2 + \ \mathbf{u}^{-\mathsf{T}}\mathbf{v}^- + \frac{\rho^-}{2}\|\mathbf{v}^-\|_2^2, \\
\text{subject to} \quad & \tilde{B}^+ = A^+Z^+, \tilde{B}^- = A^-Z^-, \\
& \mathbf{v}^+ = \text{vec}(W^+) - \text{vec}(Z^+), \mathbf{v}^- = \text{vec}(W^-) - \text{vec}(Z^-), \\
& W^+ = Z^+, W^- = Z^-, W^+ \geq 0, W^- \geq 0, \\
& \text{diag}(W^+) = 0, \text{diag}(W^-) = 0,
\end{aligned}
$$

where $\mathbf{u}^+$ and $\mathbf{u}^-$ are the Lagrange multipliers; $\rho^+, \rho^-$ ¿ 0 are the penalty parameters, and vec$(\cdot)$ is the vectorization of a matrix. The algorithm to solve the optimization problem 2 is presented in Algorithm 1.

**Algorithm 1** Learning `SLR`

**function** `SLR`$(A, \omega, \alpha, \lambda, \beta, \gamma)$
  $\rho^+ = 10, \rho^- = 10, \mathbf{u}^+_{(0)} = \mathbf{0}, \mathbf{u}^-_{(0)} = \mathbf{0}, k = 0$
  $Z^+_{(0)} = W^+_{(0)}, Z^-_{(0)} = W^-_{(0)}$
  learn $W^+_{(0)}$ and $W^-_{(0)}$ from `SLIM` (Equation 2)
  learn $\mathbf{x}_{(0)}$ and $c_{(0)}$ from `LogR` (Equation 4)
  **while** not converge **do**

$$\{W^+_{(k+1)}, W^-_{(k+1)}\} := \underset{W^+,W^-}{\text{argmin}}\ L(W^+_{(k)}, W^-_{(k)}, Z^+_{(k)}, Z^-_{(k)}, \mathbf{x}_{(k)},\ c_{(k)},\ \mathbf{u}^+_{(k)},\ \mathbf{u}^-_{(k)})$$

$$\{Z^+_{(k+1)}, Z^-_{(k+1)}\} := \underset{Z^+,Z^-}{\text{argmin}}\ L(W^+_{(k+1)}, W^-_{(k+1)}, Z^+_{(k)}, Z^-_{(k)}, \mathbf{x}_{(k)},\ c_{(k)},\ \mathbf{u}^+_{(k)},\ \mathbf{u}^-_{(k)})$$

$$\{\ \mathbf{x}_{(k+1)},\ c_{(k+1)}\} := \underset{\mathbf{x},\ c}{\text{argmin}}\ L(W^+_{(k+1)}, W^-_{(k+1)}, Z^+_{(k+1)}, Z^-_{(k+1)}, \mathbf{x}_{(k)},\ c_{(k)},\ \mathbf{u}^+_{(k)},\ \mathbf{u}^-_{(k)})$$

    $\mathbf{u}^+_{(k+1)} = \mathbf{u}^+_{(k)} + \rho^+(\text{vec}(W^+_{(k+1)} - Z^+_{(k+1)})$
    $\mathbf{u}^-_{(k+1)} = \mathbf{u}^-_{(k)} + \rho^+(\text{vec}(W^-_{(k+1)} - Z^-_{(k+1)})$
    $k = k + 1$
  **end while**
  **return** $W^+_{(k+1)}, W^-_{(k+1)}, Z^+_{(k+1)}, Z^-_{(k+1)}, \mathbf{x}_{(k+1)}$ and $c_{(k+1)}$
**end function**

In Algorithm 1, to solve for $W$, the problem boils down to a regularized least squares problem. To solve for $Z$, the problem boils down to a combination of a regularized logistic regression problem and a least squares problem. To solve for $\mathbf{x}$ and $c$, the problem boils down to a regularized logistic regression problem. All these problems can be solved by gradient descent methods. The algorithm empirically converges.

**Examples of Co-Prescribed Drugs with Similar Indications**

Table S1 presents some examples of co-prescribed drugs with similar indications.

**Additional Experimental Results**

**Table S1:** Examples of Co-Prescribed Drugs with Similar Indications

| Co-prescribed drugs | Indications |
|---|---|
| atorvastatin, lovastatin, rosuvastatin, simvastatin | high cholesterol |
| citalopram, escitalopram | depression medication |
| levofloxacin, methylprednisolone, prednisolone | arthritis, blood problems, and immune system disorders |
| fluoxetine, paroxetine, sertraline | depression and other mental illnesses |
| alitretinoin, tretinoin | acne and other skin conditions |
| conjugated estrogens, medroxyprogesterone, progesterone | birth control |

Maximum Possible Evaluation Metrics

We present the maximum possible $\max(\texttt{rec}_\texttt{P})$ and $\max(\texttt{rec}_\texttt{N})$ values that are used for $\texttt{rec}_\texttt{P}$ and $\texttt{rec}_\texttt{P}$ normalization in Table S2.

**Table S2:** Maximum Possible $\texttt{rec}_\texttt{P}$ and $\texttt{rec}_\texttt{N}$ Values

| N | $A^{\text{tst}}_{*+}$ | | | $A^{\text{tst}}_{*-}$ | | | $A^{\text{tst}}_{*u}$ | | | $A_{*all}$ | | |
|---|---|---|---|---|---|---|---|---|---|---|---|---|
| | $\texttt{rec}_\texttt{P}$ | $\texttt{rec}_\texttt{N}$ | $\text{HM}_{\text{rec}}$ | $\texttt{rec}_\texttt{P}$ | $\texttt{rec}_\texttt{N}$ | $\text{HM}_{\text{rec}}$ | $\texttt{rec}_\texttt{P}$ | $\texttt{rec}_\texttt{N}$ | $\text{HM}_{\text{rec}}$ | $\texttt{rec}_\texttt{P}$ | $\texttt{rec}_\texttt{N}$ | $\text{HM}_{\text{rec}}$ |
| 5 | 0.4567 | 0.0859 | 0.1436 | 0.9033 | 0.2824 | 0.4324 | 0.5361 | 0.0430 | 0.0804 | 0.5369 | 0.0846 | 0.1469 |
| 10 | 0.6419 | 0.1648 | 0.2596 | 0.9706 | 0.4693 | 0.6322 | 0.6739 | 0.0772 | 0.1387 | 0.6863 | 0.1517 | 0.2481 |
| 15 | 0.7440 | 0.2375 | 0.3582 | 0.9854 | 0.5951 | 0.7394 | 0.7704 | 0.1105 | 0.1932 | 0.7782 | 0.2096 | 0.3299 |
| 20 | 0.8116 | 0.3037 | 0.4412 | 0.9938 | 0.6868 | 0.8079 | 0.8347 | 0.1433 | 0.2444 | 0.8392 | 0.2611 | 0.3982 |

Parameter Study

Table S3 presents the parameter study on $\omega$ and $\alpha$ as in Equation 2 on $\texttt{SLR}$-sli for top-5 (i.e., $N = 5$) drug recommendations on $A^{\text{tst}}_{*}$. We found all 0 or very small values (e.g., $10^{-6}$) for all the other parameters $\lambda$ (parameter on $\ell_1$-norm regularization on $\texttt{SLIM}$ component), $\beta$ (parameter on $\ell_2$-norm regularization on $\texttt{LogR}$ component) and $\gamma$ (parameter on $\ell_1$-norm regularization on $\texttt{LogR}$ component) lead to optimal performance. This indicates the very minor effects of these parameters on $\texttt{SLR}$-sli. Thus, we did not present studies on these parameters.

**Table S3:** Parameter Study of $\texttt{SLR}$-sli on $A^{\text{tst}}_{*}$ ($N = 5$)

| best $\overline{\texttt{rec}_\texttt{P}}$ | | | | | | best $\overline{\texttt{rec}_\texttt{N}}$ | | | | | | best $\text{HM}_{\text{rec}}$ | | | | | |
|---|---|---|---|---|---|---|---|---|---|---|---|---|---|---|---|---|---|
| $\omega\backslash\alpha$ | 1 | 5 | 10 | 50 | 500 | $\omega\backslash\alpha$ | 1 | 10 | 20 | 50 | 100 | $\omega\backslash\alpha$ | 1 | 5 | 10 | 50 | 100 |
| 0.0005 | 0.2504 | 0.2554 | 0.2575 | 0.2505 | 0.2283 | 0.1000 | 0.3871 | 0.3941 | 0.3949 | 0.3957 | 0.3937 | 0.0010 | 0.3010 | 0.3056 | 0.3080 | 0.3040 | 0.2997 |
| 0.0050 | 0.2568 | 0.2582 | 0.2599 | 0.2517 | 0.2283 | 0.5000 | 0.3891 | 0.3950 | 0.3961 | 0.3977 | 0.3956 | 0.0050 | 0.3056 | 0.3077 | 0.3098 | 0.3049 | 0.2995 |
| 0.0100 | 0.2599 | 0.2640 | **0.2654** | 0.2572 | 0.2285 | 0.8000 | 0.3890 | 0.3944 | 0.3951 | **0.3978** | 0.3940 | 0.0100 | 0.3078 | 0.3117 | **0.3136** | 0.3091 | 0.3037 |
| 0.5000 | 0.2549 | 0.2573 | 0.2567 | 0.2454 | 0.2072 | 1.0000 | 0.3892 | 0.3943 | 0.3959 | 0.3970 | 0.3943 | 0.5000 | 0.2878 | 0.2925 | 0.2898 | 0.2812 | 0.2792 |
| 1.0000 | 0.2552 | 0.2586 | 0.2577 | 0.2464 | 0.2075 | 5.0000 | 0.3889 | 0.3935 | 0.3937 | 0.3944 | 0.3938 | 1.0000 | 0.2987 | 0.2890 | 0.2961 | 0.2739 | 0.2653 |

The best performance is **bold**.

Recall that $\omega$ is the trade-off parameter between $\texttt{SLIM}$ and $\texttt{LogR}$ components in $\texttt{SLR}$-sli, and $\alpha$ is its parameter on $\ell_2$-norm regularization on $\texttt{SLIM}$ component. Table S3 demonstrates a very similar trend in terms of $\overline{\texttt{rec}_\texttt{P}}$, $\overline{\texttt{rec}_\texttt{N}}$ and $\text{HM}_{\text{rec}}$, that is, as $\omega$ becomes larger (i.e., the weight on $\texttt{LogR}$ component becomes higher in $\texttt{SLR}$-sli), all $\overline{\texttt{rec}_\texttt{P}}$, $\overline{\texttt{rec}_\texttt{N}}$ and $\text{HM}_{\text{rec}}$ values first increase and then decrease. This demonstrates the trade-off between the $\texttt{SLIM}$ component and the $\texttt{LogR}$ component in $\texttt{SLR}$-sli. Even though, the optimal $\omega$ corresponds to a small value (e.g., 0.01). This may indicate that co-prescription pattern learning (via $\texttt{SLIM}$) is more difficult than ADR label prediction (via $\texttt{LogR}$). Similarly, as $\alpha$ becomes larger (i.e., the regularization on parameter $W$ of $\texttt{SLIM}$ becomes stronger), all $\overline{\texttt{rec}_\texttt{P}}$, $\overline{\texttt{rec}_\texttt{N}}$ and $\text{HM}_{\text{rec}}$ also first increase and then decrease. Smaller $\alpha$ will introduce larger values in $W$ compared to larger $\alpha$, and thus the relatively small optimal $\alpha$ indicates that $W$ captures strong patterns from prescriptions.

Table S4 presents parameter study of $\eta$ on $\texttt{SLR}$-sli in $A^{\text{tst}}_{*}$ for the best $\overline{\texttt{rec}_\texttt{P}}$, $\overline{\texttt{rec}_\texttt{N}}$, and $\text{HM}_{\text{rec}}$. The parameter $\eta$ is the frequency threshold for drugs in the $\texttt{SLIM}$ recommendation list (i.e., Step 2 in Section ). When higher frequency threshold is used, fewer drugs in SLIM recommendation list will be replaced, and the performance of SLR-sli in general decreases. This indicates that drug recommendation performance can be improved by considering the frequencies of drugs prescribed.

**Table S4:** Frequency Threshold Study on SLR-sli in $A_{*}^{\text{tst}}$

| $\eta$ | $\overline{\texttt{rec}_\texttt{P}}$ | $\overline{\texttt{rec}_\texttt{N}}$ | $\text{HM}_\text{rec}$ |
|---|---|---|---|
| 1 | 0.2581 | **0.3978** | 0.3120 |
| 5 | **0.2654** | 0.3880 | **0.3136** |
| 10 | 0.2626 | 0.3681 | 0.3050 |
| 15 | 0.2555 | 0.3543 | 0.2956 |
| 20 | 0.2487 | 0.3412 | 0.2866 |

The best performance is marked in **bold**.

**Table S5:** Top-$N$ Performance of SLR-sli

| N | $A_{*+}^{\text{tst}}$ | | | $A_{*-}^{\text{tst}}$ | | | $A_{*u}^{\text{tst}}$ | | | $A_{*}^{\text{tst}}$ | | |
|---|---|---|---|---|---|---|---|---|---|---|---|---|
| | $\overline{\texttt{rec}_\texttt{P}}$ | $\overline{\texttt{rec}_\texttt{N}}$ | $\text{HM}_\text{rec}$ | $\overline{\texttt{rec}_\texttt{P}}$ | $\overline{\texttt{rec}_\texttt{N}}$ | $\text{HM}_\text{rec}$ | $\overline{\texttt{rec}_\texttt{P}}$ | $\overline{\texttt{rec}_\texttt{N}}$ | $\text{HM}_\text{rec}$ | $\overline{\texttt{rec}_\texttt{P}}$ | $\overline{\texttt{rec}_\texttt{N}}$ | $\text{HM}_\text{rec}$ |
| 5 | 0.2530 | 0.4131 | 0.3137 | 0.2847 | 0.4407 | 0.3459 | 0.2541 | 0.4005 | 0.3107 | 0.2654 | 0.3836 | 0.3136 |
| 10 | 0.2956 | 0.4252 | 0.3486 | 0.3636 | 0.4408 | 0.3985 | 0.2914 | 0.4041 | 0.3385 | 0.3055 | 0.3913 | 0.3431 |
| 15 | 0.3278 | 0.4240 | 0.3697 | 0.4191 | 0.4506 | 0.4342 | 0.3061 | 0.3950 | 0.3448 | 0.3321 | 0.3940 | 0.3604 |
| 20 | 0.3538 | 0.4250 | 0.3861 | 0.4573 | 0.4631 | 0.4601 | 0.3201 | 0.3847 | 0.3493 | 0.3551 | 0.3965 | 0.3747 |

Top-$N$ Performance

Table S5 presents the SLR-sli performance with $N = 5, 10, 15$, and 20 to-avoid and safe drugs recommended when SLR-sli achieves the best $\text{HM}_\text{rec}$. As $N$ increases, all the $\text{HM}_\text{rec}$ values increase, demonstrating that more true to-avoid and safe drugs are recommended. This indicates that SLR-sli is able to rank the true to-avoid and safe drugs on top. The maximum possible $\max(\texttt{rec}_\texttt{P})$ and $\max(\texttt{rec}_\texttt{N})$ values that are used for $\overline{\texttt{rec}_\texttt{P}}$ and $\overline{\texttt{rec}_\texttt{N}}$ normalization (Equation 3 and Equation 4) is presented in Table S2.

Co-Prescription Patterns

We present the co-prescription patterns using $W^+$ and $W^-$ from SLR-sli of the best $\text{HM}_\text{rec}$ performance on $A_{*+}^{\text{tst}}$, $A_{*-}^{\text{tst}}$ and $A_{*u}^{\text{tst}}$ in Table S6, S7 and S8, respectively, and the top-10 largest values in $W^+$ and $W^-$ are presented.

**Table S6:** Co-Prescription Patterns from SLR-sli on $A_{*+}^{\text{tst}}$

| $W^+$ | | | | $W^-$ | | | |
|---|---|---|---|---|---|---|---|
| $w$ | $d_1$ | $d_2$ | freq | $w$ | $d_1$ | $d_2$ | freq |
| 0.3437 | acetaminophen | hydrocodone | 133 | 0.2033 | acetaminophen | hydrocodone | 36 |
| 0.2236 | ethinyl estradiol | etonogestrel | 38 | 0.1918 | fluticasone propionate | salmeterol | 35 |
| 0.2038 | fluticasone propionate | salmeterol | 44 | 0.1848 | **emtricitabine** | **tenofovir** | 35 |
| 0.1783 | hydrochlorothiazide | triamterene | 32 | 0.1748 | **lamivudine** | **zidovudine** | 42 |
| 0.1781 | **ezetimibe** | **simvastatin** | 68 | 0.1700 | lopinavir | **ritonavir** | 36 |
| 0.1719 | sulfamethoxazole | trimethoprim | 31 | 0.1609 | exenatide | metformin | 40 |
| 0.1672 | pamidronate | zoledronate | 42 | 0.1535 | ethinyl estradiol | norelgestromin | 23 |
| 0.1587 | acetaminophen | oxycodone | 80 | 0.1466 | acetylsalicylic acid | clopidogrel | 29 |
| 0.1471 | salbutamol | salmeterol | 37 | 0.1228 | **ritonavir** | **tenofovir** | 36 |
| 0.1419 | acetylsalicylic acid | ramipril | 65 | 0.1142 | **atazanavir** | **ritonavir** | 25 |

In this table, "$w$" is the value of the entry corresponding to the drug pairs in $W^+$/$W^-$; "$d_1$" and "$d_2$" are the two drugs in the drug pair, and "freq" is the frequency of the corresponding drug pairs in training data $A^+$ and $A^-$. Drugs that are reported in SIDER to induce myopathy on their own are **bold**.

Case Study

**Table S7:** Co-Prescription Patterns from SLR-sli on $A^{\text{tst}}_{*\text{-}}$

| $W^+$ | | | | $W^-$ | | | |
|---|---|---|---|---|---|---|---|
| $w$ | $d_1$ | $d_2$ | freq | $w$ | $d_1$ | $d_2$ | freq |
| 0.5659 | ethinyl estradiol | etonogestrel | 38 | 0.4854 | acetaminophen | hydrocodone | 36 |
| 0.5309 | acetaminophen | hydrocodone | 13 | 0.4660 | ethinyl estradiol | norelgestromin | 23 |
| 0.4662 | hydrochlorothiazide | triamterene | 32 | 0.4625 | **emtricitabine** | **tenofovir** | 35 |
| 0.4541 | sulfamethoxazole | trimethoprim | 31 | 0.4519 | fluticasone propionate | salmeterol | 35 |
| 0.4430 | fluticasone propionate | salmeterol | 44 | 0.4039 | lopinavir | **ritonavir** | 36 |
| 0.3633 | acetaminophen | codeine | 27 | 0.3393 | budesonide | formoterol | 17 |
| 0.3607 | amoxicillin | clavulanate | 22 | 0.3324 | conjugated estrogens | medroxyprogesterone | 17 |
| 0.3596 | pamidronate | zoledronate | 42 | 0.3311 | fluorouracil | leucovorin | 18 |
| 0.3540 | carbidopa | levodopa | 17 | 0.3278 | **lamivudine** | **zidovudine** | 42 |
| 0.3345 | **emtricitabine** | **tenofovir** | 22 | 0.2974 | acetylsalicylic acid | clopidogrel | 29 |

In this table, "$w$" is the value of the entry corresponding to the drug pairs in $W^+$/$W^-$; "$d_1$" and "$d_2$" are the two drugs in the drug pair, and "freq" is the frequency of the corresponding drug pairs in training data $A^+$ and $A^-$. Drugs that are reported in SIDER to induce myopathy on their own are **bold**.

**Table S8:** Co-Prescription Patterns from SLR-sli on $A^{\text{tst}}_{*\text{u}}$

| $W^+$ | | | | $W^-$ | | | |
|---|---|---|---|---|---|---|---|
| $w$ | $d_1$ | $d_2$ | freq | $w$ | $d_1$ | $d_2$ | freq |
| 0.8906 | carbidopa | levodopa | 17 | 0.9154 | ethinyl estradiol | norelgestromin | 23 |
| 0.8747 | ethinyl estradiol | etonogestrel | 38 | 0.8934 | amphetamine | dextroamphetamine | 8 |
| 0.8348 | amoxicillin | clavulanate | 22 | 0.8851 | buprenorphine | naloxone | 9 |
| 0.8080 | sulfamethoxazole | trimethoprim | 31 | 0.8750 | sulfamethoxazole | trimethoprim | 8 |
| 0.7979 | buprenorphine | naloxone | 9 | 0.8276 | **emtricitabine** | **tenofovir** | 35 |
| 0.7973 | ethinyl estradiol | norgestimate | 10 | 0.7385 | piperacillin | tazobactam | 3 |
| 0.7755 | prazepam | **venlafaxine** | 4 | 0.7309 | lopinavir | **ritonavir** | 36 |
| 0.7734 | hydrochlorothiazide | triamterene | 32 | 0.7160 | fluorouracil | leucovorin | 18 |
| 0.7400 | cyproterone | **simvastatin** | 4 | 0.7158 | acetaminophen | hydrocodone | 36 |
| 0.7389 | amphetamine | dextroamphetamine | 6 | 0.7126 | budesonide | formoterol | 17 |

In this table, "$w$" is the value of the entry corresponding to the drug pairs in $W^+$/$W^-$; "$d_1$" and "$d_2$" are the two drugs in the drug pair, and "freq" is the frequency of the corresponding drug pairs in training data $A^+$ and $A^-$. Drugs that are reported in SIDER to induce myopathy on their own are **bold**.

Table S9 presents some examples of testing prescriptions and their recommended to-avoid drugs from SLR-sli such that the corresponding new prescriptions (i.e., testing prescriptions and recommended drugs together) are ADR-inducing.

Table S10 presents some examples of testing prescriptions and their recommended safe drugs from SLR-sli such that the corresponding new prescriptions (i.e., testing prescriptions and recommended drugs together) are ADR-free.

**Table S9:** To-Avoid Drug Recommendation from SLR-sli

| subset | testing prescription | recommendation |
| --- | --- | --- |
| $A^{tst}_{*+}$ | calcipotriol, **cerivastatin**, **fenofibrate**, gliclazide, **rosuvastatin**, sulfasalazine | **atorvastatin** |
| | acetylsalicylic acid, amlodipine, bisoprolol, felodipine, finasteride, lisinopril | **atorvastatin** |
| | acetylsalicylic acid, clopidogrel, cyclosporine, **fluvastatin**, gabapentin, mycophenolate mofetil, pantoprazole, prednisone | metoprolol |
| | carboplatin, cetuximab, fluticasone propionate, folic acid, hydroxocobalamin, pemetrexed, salbutamol | salmeterol |
| | acetylsalicylic acid, atenolol | **simvastatin** |
| | fusidic acid, **simvastatin** | ramipril |
| | propofol, valproic acid | lamotrigine |
| $A^{tst}_{*-}$ | acetylsalicylic acid, bisoprolol, furosemide, metformin, ramipril | **atorvastatin** |
| | acetylsalicylic acid, amlodipine, **atorvastatin**, isosorbide mononitrate, lisinopril, olmesartan, omeprazole, quinine | atenolol |
| | dicyclomine, gabapentin, lansoprazole, lorazepam, pamidronate, zoledronate, zolpidem | oxycodone |
| | amlodipine, **ezetimibe** | **simvastatin** |
| | irbesartan, metformin | **rosuvastatin** |
| | **dexamethasone**, folic acid | lenalidomide |
| | clonazepam, phenobarbital | carbamazepine |
| $A^{tst}_{*u}$ | acetaminophen, darunavir, **dexamethasone**, esomeprazole, ondansetron, **ritonavir**, sulfamethoxazole, **tenofovir**, trabectedin, trimethoprim | **emtricitabine** |
| | acetylsalicylic acid, carvedilol, isosorbide mononitrate, nitroglycerin, ramipril, ranolazine, trimethoprim | **simvastatin** |
| | acetylsalicylic acid, alprazolam, **fluoxetine**, hydrocodone, **phenytoin**, **rosuvastatin** | acetaminophen |
| | acetylsalicylic acid, amlodipine, esomeprazole, hydrochlorothiazide, potassium chloride, sorafenib, triamterene | furosemide |
| | **dexamethasone**, triazolam | **atorvastatin** |
| | zuclopenthixol | **simvastatin** |
| | mycophenolate mofetil, **simvastatin** | cyclosporine |
| | risperidone | haloperidol |
| $A^{tst}_{*}$ | acetylsalicylic acid, buprenorphine, clopidogrel, flucloxacillin, fusidic acid, metronidazole, **pregabalin**, ramipril | **atorvastatin** |
| | acetylsalicylic acid, amlodipine, bisoprolol, felodipine, finasteride, lisinopril | **atorvastatin** |
| | acetaminophen, amitriptyline, bupropion, **fluoxetine**, gabapentin, oxycodone, sildenafil, **simvastatin**, valdecoxib, **venlafaxine** | hydrocodone |
| | acetaminophen, cephalexin, diphenhydramine, etonogestrel, levothyroxine, methimazole, metoprolol, nadolol, salbutamol | ethinyl estradiol |
| | **fluoxetine** | **paroxetine** |
| | prazepam | **atorvastatin** |
| | **clarithromycin**, **simvastatin** | amoxicillin |
| | cyclosporine, methotrexate | mycophenolate mofetil |

In this table, "recommendation" represents the recommended drug.Drugs that are reported in SIDER to induce myopathy on their own are **bold**.

**Table S10:** Safe Drug Recommendation from SLR-sli

| subset | testing prescription | recommendation |
|---|---|---|
| $A^{tst}_{*+}$ | abacavir, **atazanavir**, **lamivudine**, lopinavir, **ritonavir** | **zidovudine** |
| | cyclophosphamide, doxorubicin, prednisolone, vincristine | **dexamethasone** |
| | cyclophosphamide, cytarabine, **dexamethasone**, doxorubicin, methotrexate, thioguanine | vincristine |
| | busulfan, cyclophosphamide, cyclosporine, methotrexate | prednisolone |
| | acetylsalicylic acid, **dexamethasone** | **bortezomib** |
| | acetylsalicylic acid, varenicline | **simvastatin** |
| | **methylprednisolone**, prednisolone | azathioprine |
| | carboplatin, vinorelbine | cetuximab |
| $A^{tst}_{*-}$ | darunavir, **emtricitabine**, etravirine, **ritonavir**, **tenofovir**, tipranavir | **raltegravir** |
| | cyclophosphamide, cytarabine, doxorubicin, methotrexate, thioguanine, vincristine | **dexamethasone** |
| | **bortezomib**, cisplatin, cyclophosphamide, **dexamethasone**, etoposide, thalidomide | melphalan |
| | bleomycin, cyclophosphamide, doxorubicin, etoposide, procarbazine, vincristine | prednisone |
| | stavudine, **tenofovir** | **emtricitabine** |
| | sertraline, topiramate | **phenytoin** |
| | **atorvastatin**, solifenacin | amlodipine |
| | everolimus, prednisolone | mycophenolic acid |
| $A^{tst}_{*u}$ | **efavirenz**, **lamivudine**, lopinavir, **ritonavir**, **tenofovir**, **zidovudine** | **emtricitabine** |
| | carboplatin, diphenhydramine, granisetron, paclitaxel, ranitidine | **dexamethasone** |
| | **bortezomib**, cisplatin, cyclophosphamide, **dexamethasone**, etoposide, melphalan | doxorubicin |
| | bleomycin, cyclophosphamide, doxorubicin, etoposide, prednisolone, procarbazine | vincristine |
| | **emtricitabine** , fosamprenavir | **ritonavir** |
| | albendazole | **dexamethasone** |
| | **hydroxychloroquine** , risedronate | prednisone‘ |
| | calcium | valproic acid |
| $A^{tst}_{*}$ | lopinavir, nevirapine, **raltegravir**, **ritonavir**, **tenofovir**, **zidovudine** | **lamivudine** |
| | cyclophosphamide, cytarabine, doxorubicin, methotrexate, thioguanine, vincristine | **dexamethasone** |
| | **atazanavir**, **efavirenz**, **emtricitabine**, **lamivudine**, **tenofovir**, **zidovudine** | stavudine |
| | acetylsalicylic acid, ciprofloxacin, erythromycin, meropenem, metronidazole, teicoplanin, warfarin | clopidogrel |
| | nevirapine, **tenofovir** | **zidovudine** |
| | estradiol, levothyroxine | **progesterone** |
| | **rofecoxib**, rosiglitazone | glyburide |
| | amlodipine, olmesartan | acetylsalicylic acid |

In this table, “recommendation” represents the recommended drug. Drugs that are reported in SIDER to induce myopathy on their own are **bold**.
